# Dopamine synchronizes hippocampal-prefrontal networks

**DOI:** 10.1101/2023.06.07.544112

**Authors:** Benedito A de Oliveira-Junior, Fernando E Padovan-Neto, Nandakumar S Narayanan, Joao P Leite, Rafael N Ruggiero

## Abstract

Interactions among brain networks support cognition and behavior. Oscillatory activity may contribute to this process by coordinating distant neuronal populations. A crucial step toward elucidating brain function and its emergent properties is understanding the mechanisms that dynamically modulate functional connectivity in response to environmental challenges and physiological demands. Dopamine (DA) is well-positioned to modulate this process, given its role in altering neuronal excitability and regulating oscillatory cognitive networks, such as the hippocampus-prefrontal cortex (HPC-PFC) networks. However, it is mostly unknown how dopamine modulates HPC-PFC synchrony. Here, we recorded HPC and PFC activity in rats while manipulating dopamine levels. Our results show that dopamine dose-dependently induces HPC-PFC theta synchrony. This effect was not reproduced by selective activation of D_1_-like (SKF-38393) or D_2_-like (quinpirole) receptors, nor by the D_1_/D_2_ non-selective agonist apomorphine. Thus, dopamine’s modulatory effects on HPC-PFC activity may depend on the precise interplay between dopamine receptor subtypes. Our findings provide evidence that dopaminergic neurotransmission plays a central role in regulating HPC-PFC oscillatory dynamics, specifically promoting synchronization in slow-frequency oscillations.

## Introduction

Brain activity underlying behavior is highly flexible and emerges from the coordinated dynamics of distributed neuronal networks. Neural oscillations are believed to promote the formation of neuronal ensembles and facilitate communication between ensembles from distinct brain regions (Fries, 2005; Buzsáki, 2010). Oscillations are possibly the most energy-efficient mechanism for the temporal coordination of neural activity (Buzsáki & Draguhn, 2004). Thus, investigating how oscillatory synchrony operates is particularly interesting for understanding the dynamic coordination of brain activity.

Fries (2005; 2015) proposes that interareal coherence creates brief, milliseconds-scale windows in which inputs from one neuronal group are preferentially transmitted and read out by another. Because coherence is flexible and state-dependent, it may be tuned by neurophysiological processes such as synaptic plasticity (Uhlhaas et al., 2010) and neuromodulation, particularly dopaminergic (Beeler & Dreyer, 2019), thereby enabling selective, dynamic routing of information across brain networks.

Dopamine (DA), acting through D_1_- and D_2_-like receptors, powerfully influences neuronal excitability and synaptic efficacy (Goto & Grace, 2005; Lezcano & Bergson, 2002; Robinson & Sohal, 2017), which could provide the basis for DA to contribute to interareal synchronization. Specifically, the participation of DA in oscillatory synchrony has been investigated in the hippocampus-prefrontal cortex (HPC-PFC) networks, which are central to cognition and emotion and are coordinated by hippocampal theta rhythms (Benchenane et al., 2011; Ruggiero et al., 2021; Sigurdsson & Duvarci, 2016; Spellman et al., 2015). In working memory tasks, HPC-PFC theta synchrony varies dynamically and is usually correlated to individual performance (Benchenane et al., 2011; Fujisawa & Buzsáki, 2011; Sigurdsson et al., 2010; O’Neill et al., 2013). Interestingly, it is known that DA release in the cortex increases during working memory tasks (Phillips et al., 2004). Notably, Benchenane et al. (2010) showed that HPC-PFC theta coherence could be reproduced by PFC local DA administration in anesthetized rats. Furthermore, both agonism and antagonism of D_1_- and D_2_-like DA receptors are shown to be associated with changes in HPC-PFC theta synchrony (Xu et al., 2016; Perreault et al., 2017; Gener et al., 2019), reinforcing the DA proposal as a neural synchrony modulator (Beeler & Dreyer, 2019).

Together, the literature suggests that dopaminergic neurotransmission plays a role in regulating HPC-PFC oscillatory activity, influencing power or coherence (Benchenane et al., 2010; Xu et al., 2016; Perreault et al., 2017; Gener et al., 2019). However, this process is still poorly understood. For instance, in a small sample, Benchenane et al. (2010) reported that local DA administration in the PFC increased theta coherence without altering theta power in either PFC or HPC. Since coherence is a measure dependent on power, these findings suggest that there may also be phase synchrony modulation, i.e., a power-unbiased and volume-conduction-free synchronization. In this study, we tested the hypothesis that DA modulates HPC-PFC oscillatory phase synchrony. Here, we investigated the impact of DA administration on HPC-PFC functional connectivity, and whether these effects are reproduced by selective D_1_- and D_2_-like DA single-receptor agonism or non-specific apomorphine (APO) agonism. Our findings reveal that DA induces HPC-PFC theta phase synchrony dose-dependently, and receptor agonists do not fully reproduce this effect. Therefore, our data suggest that DA has an essential but complex role in regulating HPC-PFC oscillatory activity, influencing not only power and coherence but also phase synchrony in theta frequencies.

## Materials and Methods

### Animals

Forty-two male seven-week-old Sprague Dawley rats were used in this study. The animals were housed in standard rodent cages and kept in a controlled-temperature room (24 ± 2 °C) that operated in cycles of 12 hours of light and 12 hours of dark, with lights on at 7:00 a.m. All animals had ad libitum access to food and water. This study was approved by the local ethics committee on animal experimentation (Ribeirão Preto Medical School, University of São Paulo; protocol number: 74/2021).

### Surgery and electrode implantation

Rats were anesthetized under urethane (1.2 g/Kg in NaCl 0.15 M, i.p., Sigma-Aldrich) and placed in a stereotaxic frame (Kopf Instruments, EUA) equipped with a heating pad (Insight, Brazil) for temperature maintenance (37 ± 0.5 °C). Once the skull was exposed, burr holes were drilled, targeted at the left medial PFC (anteroposterior, AP: + 3.2 mm; medial-lateral, ML: - 0.5 mm; dorsal-ventral, DV: - 2.9 mm) and the left dorsal HPC (AP: - 5.0 mm, ML: - 2.7 mm, DV: - 3.1 mm) for local field potential (LFP) recordings, and the right lateral ventricle (LV; AP: - 0.5 mm, ML: - 1.8 mm, DV: - 3.5 mm) for cannula implantation to deliver intracerebroventricular (i.c.v.) injections (Fig. 1A).

**Figure 1.**
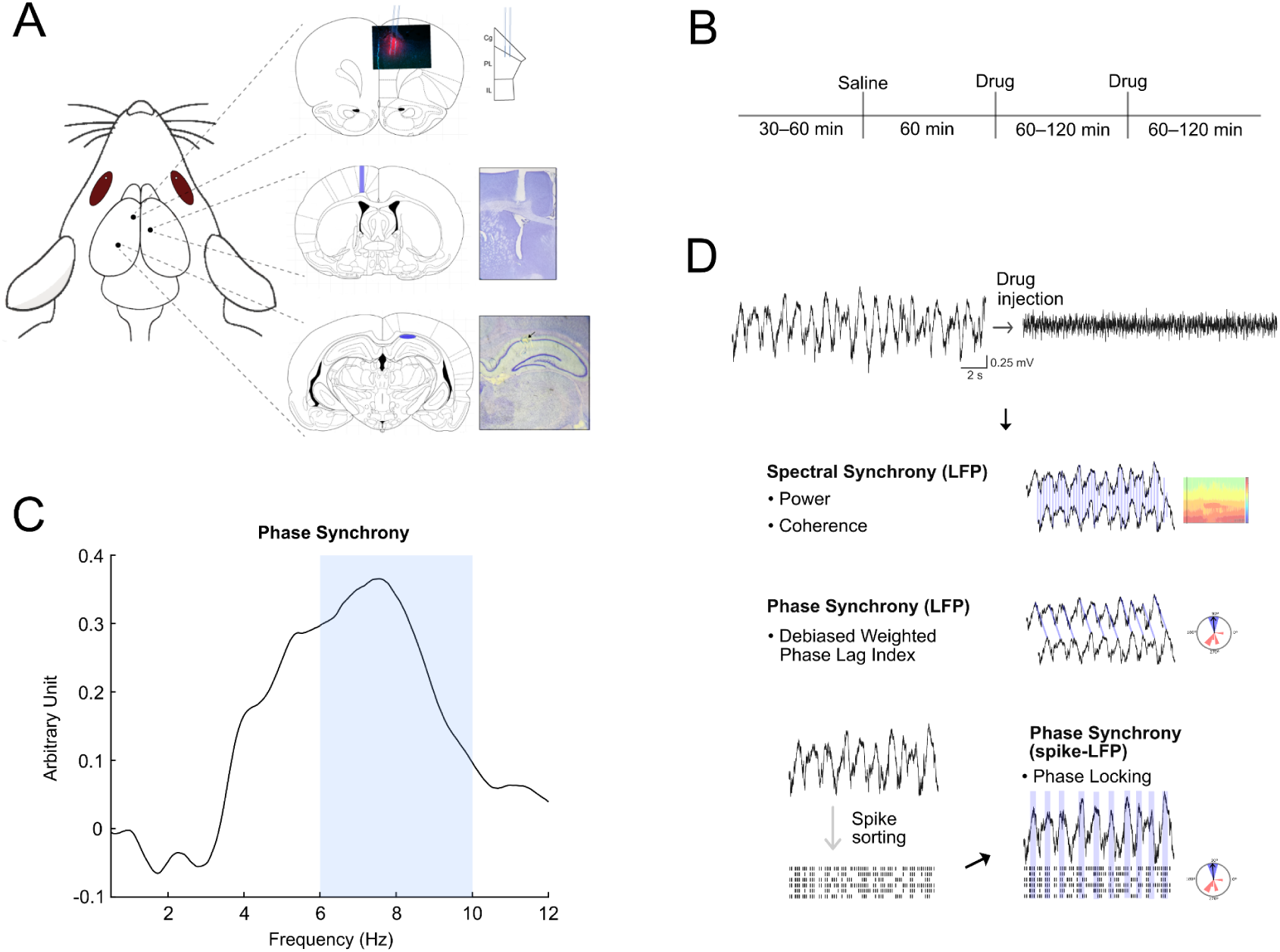
Experimental design and analytical workflow. (A) Surgical targeting and post hoc histology. Left: schematic showing hippocampal (HPC) and medial prefrontal cortex (PFC) recording locations from forty-two recordings, and a single intracerebroventricular (i.c.v.) cannula for drug injection. Right, top-to-bottom: representative coronal sections showing a 32-channel silicon probe spanning PFC (left hemisphere), a cannula tract into the right lateral ventricle for drug infusion, and an electrolytic lesion in CA1 (left hemisphere) marking tungsten-wire electrode placement for recordings. (B) Recording timeline. Animals underwent a 30-60 min baseline, followed by post-injection epochs (saline, then one or two drug doses; order counterbalanced). (C) The 6-10 Hz range was selected for LFP power and HPC-PFC connectivity analyses since it encompassed the peak of the post-minus-pre difference in phase synchrony (dwPLI). As dwPLI selectively captures non-zero phase-lag interactions, this approach minimizes the contribution of volume-conducted signals. (D) Analysis pipeline. Top: representative raw PFC LFP trace before and after drug infusion. Middle: spectral synchrony (power spectral density and magnitude-squared coherence) and phase-synchrony (debiased weighted phase-lag index; dwPLI) measures computed from HPC-PFC LFPs. Bottom: spike sorting followed by spike-LFP synchrony (phase locking) analysis.

A single stereotrode (two intertwined, 50-µm, Teflon-coated tungsten wires) or a 64-channel acute silicon probe (E64+R-50-S4-L6-200 NT, ATLAS Neuroengineering, Belgium) was implanted in the PFC, and a single stereotrode was implanted in the HPC. Each stereotrode was attached to a silver wire soldered into an 8-channel connector (Omnectics Connector Corporation, EUA). Silicon probes and Omnetics connectors were coupled to 32-channel Intan headstage preamplifiers (Intan Technologies, EUA).

### Extracellular recordings

The recordings were acquired using the Open Ephys acquisition system (Siegle et al., 2017). For recordings with silicon probes, the following parameters were used for signal acquisition: 1000x gain, 30 kHz sampling rate, 0.5-7603.8 Hz bandpass filtering. For recordings with tungsten wire stereotrodes, signals were acquired with 1000x gain, a 5 kHz sampling rate, and 0.1-1000 Hz bandpass filtering.

### Drugs and experimental design

Apomorphine hydrochloride, SKF-38393 hydrochloride, and quinpirole hydrochloride (Sigma-Aldrich, USA) were dissolved in sterile 0.9% NaCl. Dopamine hydrochloride (Sigma-Aldrich, USA) was dissolved in 0.04 mg/mL ascorbic acid to prevent oxidation. Aliquots were stored at −80 °C, and each aliquot was thawed once on the day of use. The doses were as follows: apomorphine 0.75, 1.5, or 3 mg/kg (i.p.); dopamine 100 or 500 nmol (i.c.v.); SKF-38393 1 or 10 µg (i.c.v.); quinpirole 1 or 10 µg (i.c.v.). All drugs except apomorphine were infused at a rate of 0.5 μL/min for a total volume of 1µL using a Hamilton microsyringe (10 μL) connected to polyethylene tubing and a 30-gauge cannula. Each animal was given a saline injection followed by a drug injection. The interval between injections was at least 60 minutes to minimize the influence of the previous drug on the next injection (Fig. 1B).

### Brain extraction and tissue processing

At the end of the recordings, electrolytic lesions were made to mark the electrode insertion sites. Rats were then euthanized with an additional dose of thiopental (25 mg/kg, i.p.), and brains were rapidly removed. Tissue was post-fixed in 4% paraformaldehyde (PFA) for 12 h and cryoprotected in 30% sucrose for 72 h. Semi-frozen coronal sections were then obtained, stained with hematoxylin and eosin, and examined to verify electrode tracks and insertion sites.

### Data analysis

Signal analyses were performed using custom codes in MATLAB software (Mathworks, USA). LFP signals were anti-alias filtered and downsampled to 500 Hz using MATLAB’s *decimate* function before spectral and connectivity analyses. LFP power and interareal connectivity were analyzed for delta (0.5-3Hz) and theta (6-10 Hz) bands. This theta band range was selected because it encompassed the peak of the drug-related change in HPC-PFC phase synchrony and therefore best captured the interareal theta effect examined in our study (Fig. 1C).

### Spectral power

Power spectral density (PSD) was calculated using Welch’s method via the Fast Fourier Transform algorithm available in the MATLAB software. We used 3s time windows with 50% overlap and 2^16^ FFT points. We averaged the PSD estimates across trials and animals, respectively. The relative power was calculated by dividing the PSD estimate by the integrated power over the frequency range of 0.5-55 Hz. Whole-recording spectrograms and coherograms were computed with the Chronux toolbox (Bokil et al., 2010) using a 5 s moving window advanced in 2.5 s steps, a multitaper method (time-bandwidth product = 3, 5 tapers), a sampling rate of 500 Hz, and a frequency range of 0.5-55 Hz.

### Functional connectivity

Spectral coherence and debiased weighted phase lag index (dwPLI - used here to evaluate oscillatory phase synchrony; Vinck et al., 2011) measures were estimated using cross PSDs of HPC and PFC calculated using modified Welch’s periodogram method. Spectral coherence is the same as that obtained by MATLAB’s mscohere function and indicates how well the signals being compared match each other at each frequency. dwPLI was calculated as described by Vinck et al. (2011) and quantifies the phase relationship between oscillations, thereby reducing zero-lag contamination due to volume conduction. For coherograms of the whole recording, we used the cohgramc function from the Chronux toolbox (Bokil et al., 2010) with the multi-tapered window method and the same parameters described for spectrograms (Fig. 1D).

### Spike sorting and single-unit analysis

Spike sorting was performed using *Kilosort 2.5* (Pachitariu et al., 2021). Quality control of individual units was performed manually using the *phy* interactive interface (https://github.com/cortex-lab/phy/). Firing-rate and spike-LFP phase-locking analyses were performed using custom MATLAB scripts. Mean firing rate was quantified for baseline and post-injection epochs, and spike-LFP phase locking was estimated as the phase-locking value between PFC spike times and the phase of the simultaneously recorded local PFC LFP. For spike-LFP analyses, the LFP was filtered in the 4-10 Hz band to provide more stable phase estimates at the single-unit level. Statistical analyses were based on 4-minute windows immediately before and after injection (Fig. 1D).

### Statistical analysis

We used the Shapiro-Wilk normality test to verify the data distribution. To compare two experimental groups or two conditions, we used the paired or unpaired two-tailed Student’s t-test, or the non-parametric Wilcoxon matched-pairs signed-rank test using the GraphPad Prism 9.0 software (GraphPad Software, EUA).

All effects were tested using linear mixed-effects models (LMM). For DA, we fitted an LMM including the fixed factors Condition (Sal vs. DA) and DoseGroup (100 vs. 500 nmol), with a random intercept for each subject (animal nested within dose). For APO, SKF, and QUIN, we modeled Dose (saline plus the respective drug dose) as a fixed factor, with random intercepts for animal and, when applicable, an additional random intercept for injection order. Post hoc comparisons were conducted using estimated marginal means (EMMs) with a Holm adjustment for multiple comparisons. The models were implemented in R (v4.5.2; R Core Team, 2025) and run in Google Colab (Google, n.d.), a hosted Jupyter notebook environment (Kluyver et al., 2016).

## Data and Code Availability

Electrophysiological data recorded for this study are publicly available at https://dandiarchive.org/dandiset/001704/draft. The code for the analysis and figure generation is accessible at https://github.com/oliveirajun/Oliveira-Junior_2026_DA.

## Results

### DA increases HPC-PFC theta functional connectivity and PFC spike-LFP synchrony

We first investigated the effects of dopamine administration on HPC-PFC functional activity. Relative to baseline, i.c.v. injection of 100 nmol of DA did not induce reliable changes of theta relative power (Fig. 2A), although the PFC theta/delta relative power ratio shows theta predominance after the drug injection (Fig. 2C). At 500 nmol, DA induced a general increase in theta relative power (PFC: theta, t_(12)_=2.8, p=0.01; HPC: theta, t_(12)_=2.7, p=0.02; n_rats_=13; Fig. 2B). Consistent with these statistics, the linear mixed model reveals a significant main effect for theta power in both PFC and HPC (DA vs. Sal - PFC: F(_1, 21_)=8.97, p<0.01; HPC: F(_1, 42_)=5.96, p=0.02, n_rats_=13; Fig 2D) for the highest dose (DA 500 nmol Holm-adjusted: PFC, p<0.01; HPC, p=0.01).

**Figure 2.**
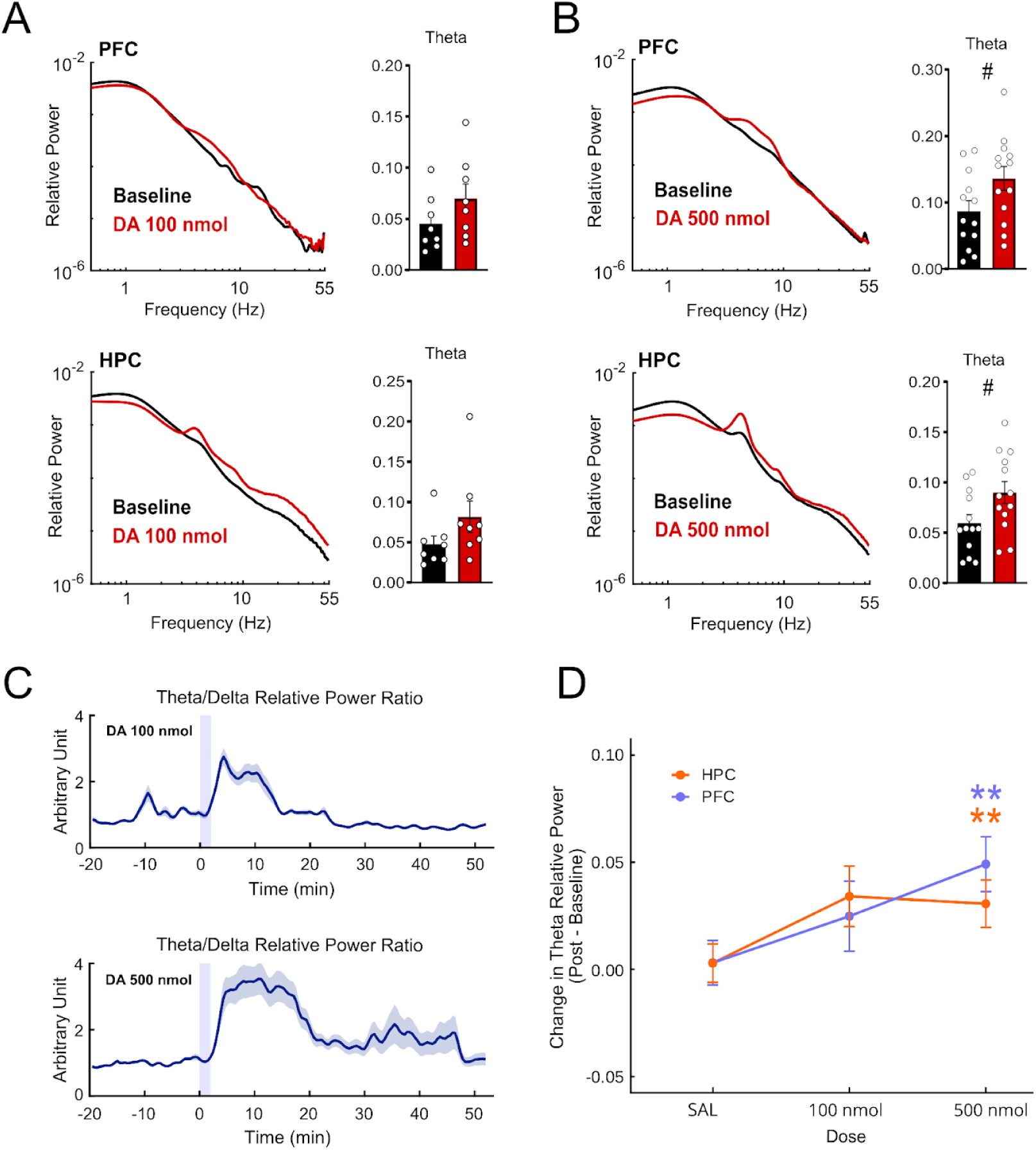
Dopamine (DA) increases theta relative power in PFC and HPC in a dose-dependent manner. (A-B) Log-log power spectra (10-min post-injection vs. baseline) for PFC (top) and HPC (bottom) after DA 100 nmol (A) and 500 nmol (B). Bar plots show the integrated relative power in the theta band (6-10 Hz). DA 500 nmol increased theta power in both regions (n_rats_=13), with smaller, nonsignificant trends at 100 nmol (n_rats_=8); paired two-tailed t-test; error bars represent SEM; (C) Time course of PFC theta/delta relative power ratio (normalized to baseline; shaded bands represent SEM; vertical shaded bar represents Injection window). DA induces an early increase in the theta/delta relative power, which is prolonged at the highest dose. (D) Dose-response of the change in theta relative power (post - baseline) for PFC and HPC. Points and error bars are linear mixed model (LMM) estimated marginal means ± SEM. # p<0.05 for t-tests; * p<0.05 for LMM main effect; ** p<0.05 for LMM Holm-adjusted post hoc pairwise comparison.

Similarly, interareal synchrony effects were more pronounced following the highest DA dose. While DA 100 nmol did not change the HPC-PFC oscillatory synchrony (Fig. 3A, C), DA 500 nmol reliably increased coherence at theta (t_(12)_=2.7, p=0.02, n_rats_=13; Fig. 3B, top), as well as phase synchrony (dwPLI; theta, t_(12)_=3.8, p<0.01, n_rats_=13; Fig. 3B, bottom). This effect was also observed when comparing the effects of DA 500 nmol with saline on theta coherence (DA vs. Sal - main effect: F_(1, 21)_=4.9, p=0.04) and theta phase synchrony (DA vs. Sal: F_(1, 21)_=16.28, p<0.001; DA 500 nmol Holm-adjusted: coherence, p=0.02; phase synchrony, p<0.001; Fig. 3E). Accordingly, DA 500 nmol showed a significantly stronger theta modulation compared to DA 100 nmol (F_(1, 21)_=4.62, p=0.04). The dose-response curve illustrates this difference, highlighting the stronger modulation of phase synchrony, while showing a smaller overall dose effect over coherence (Fig. 3E).

**Figure 3.**
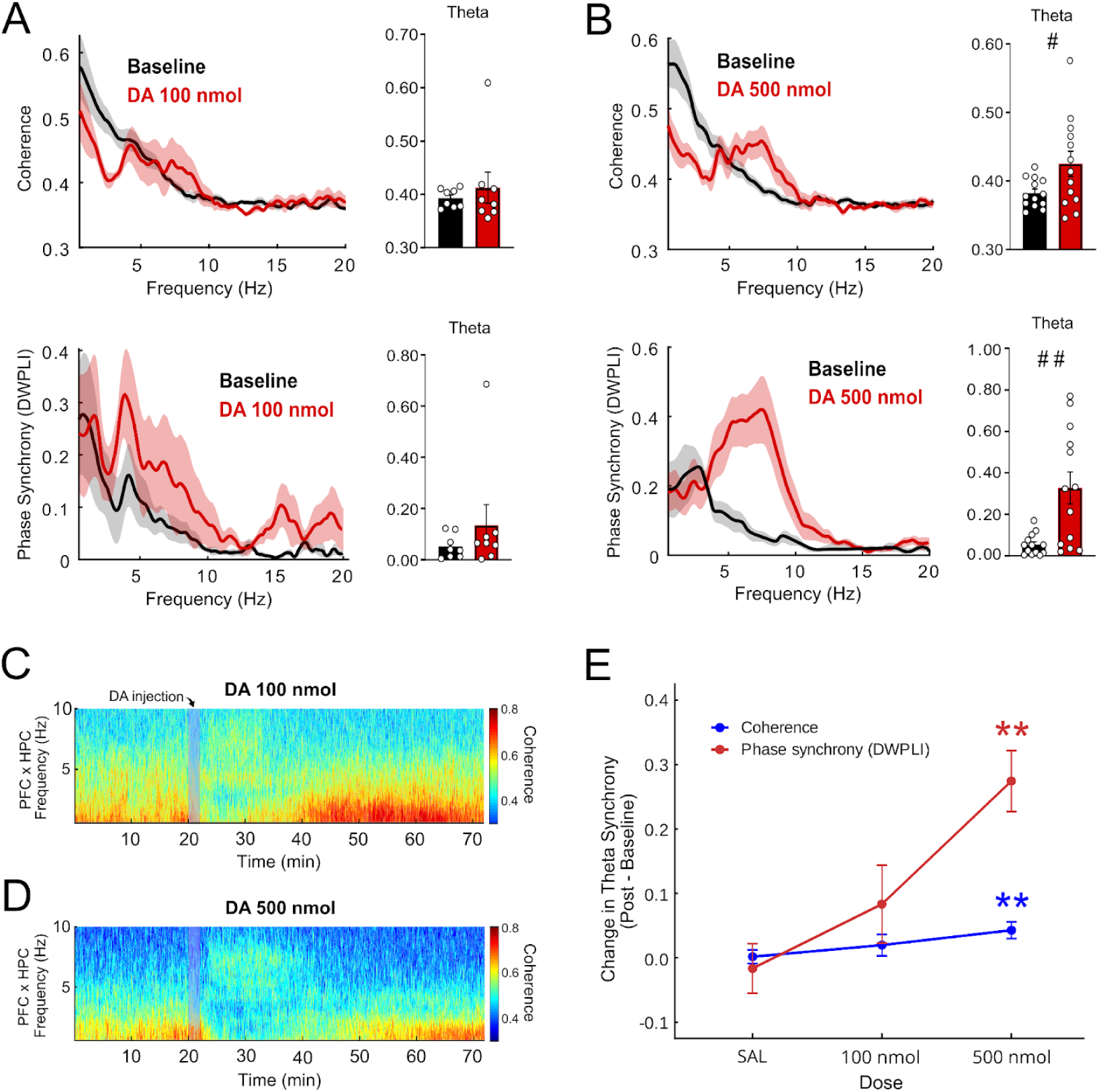
DA induces HPC-PFC theta phase synchrony and coherence in a dose-dependent manner. (A-B) Spectral coherence (top) and phase synchrony by debiased weighted phase-lag index (dwPLI; bottom) averaged over 10-min post-injection vs. baseline after DA 100 nmol (A) and 500 nmol (B). DA 500 nmol induces theta coherence and phase synchrony (n_rats_=13), while DA 100 nmol showed weaker increases (n_rats_=8); paired two-tailed t-tests. Error bars represent SEM. (C-D) HPC-PFC coherograms for each dose. The vertical bar represents the 2-minute injection period. (E) Dose-response of change in theta synchrony. DA 500 nmol shows higher theta coherence and phase synchrony modulation compared to DA 100 nmol and saline. Points and error bars are linear mixed model (LMM) estimated marginal means ± SEM. # p<0.05 and ## p<0.01 for t-tests statistics; ** p<0.05 for LMM Holm-adjusted post hoc pairwise comparison.

To evaluate whether the DA modulatory effect is reflected in the dynamics between neuronal and oscillatory activities in the PFC, we examined the phase locking of PFC single-unit activity to local theta oscillations after DA 500 nmol administration. DA induced a reduction in firing rate (t_(130)_=2.73, p<0.01, n_neurons_=131, n_rats_=6; Fig. 4A). Phase locking analysis revealed an increase in spike-LFP synchrony (t_(48)_=-2.47, p=0.02, n_neurons_=49, n_rats_=6; Fig. 4B, right) together with a change in the preferred firing phase relative to theta cycles (Fig. 4B, left), which did not occur after saline injection (Fig. 4C-D). These data are consistent with reports that DA receptor-dependent 4 Hz oscillations in PFC exhibit strong spike-field coherence and depend on intact dopaminergic signaling (Parker et al., 2014; 2017; Kim et al., 2017; 2019). Together, LFP and spike-LFP results indicate that DA modulates the dynamics of HPC-PFC activity dose-dependently, altering the preferential phase of neuronal firing and inducing spike-LFP theta synchrony in the PFC, resulting in increased functional connectivity in the pathway, particularly at the highest dose applied.

**Figure 4.**
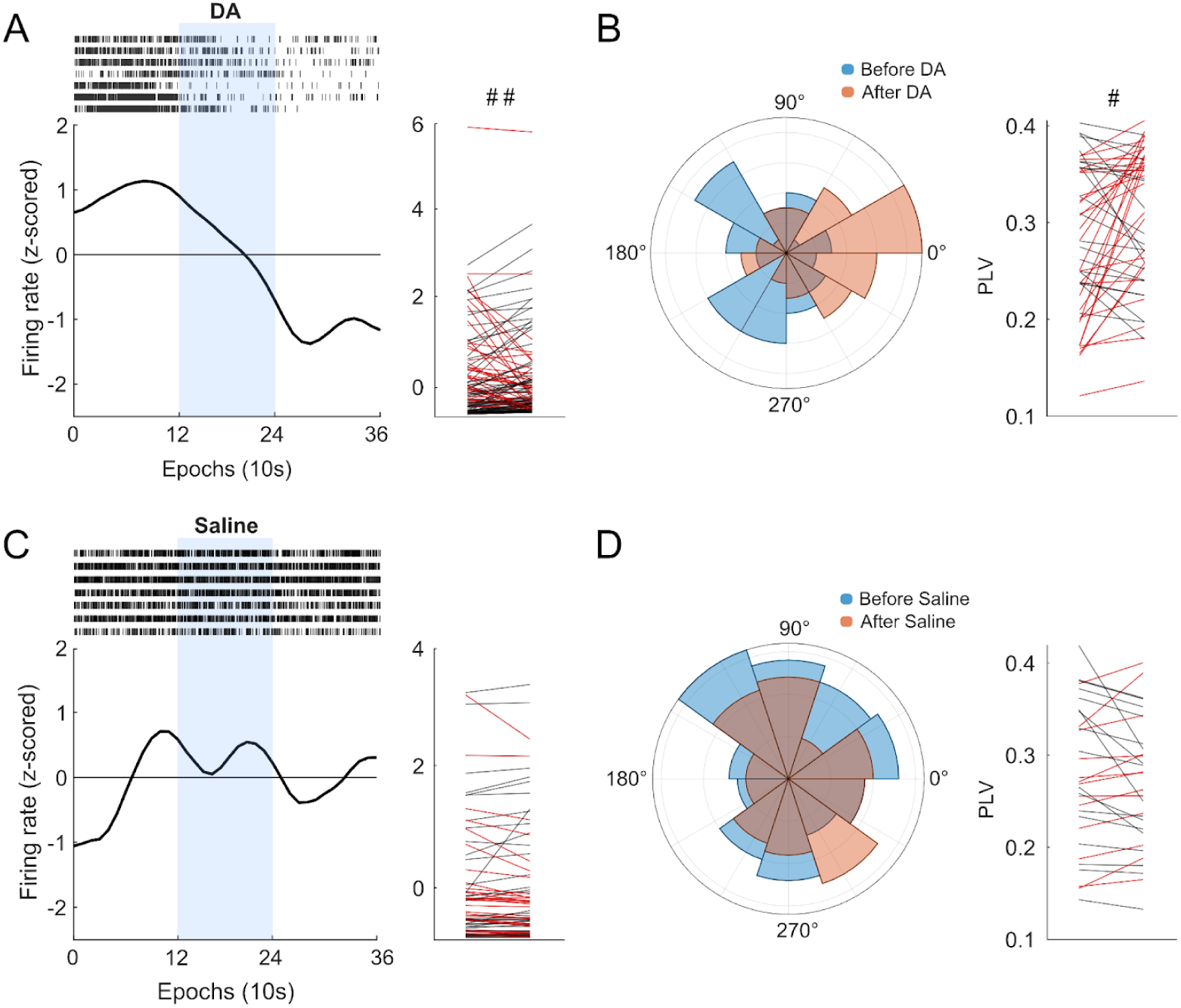
DA reorganizes PFC single-unit activity and enhances spike-LFP theta phase locking. (A) DA 500 nmol reduces PFC neuronal firing rates. Spike raster (top) displays spikes from representative neurons. Population z-scored firing-rate histogram shows the average decrease in neuronal activity after DA injection (n_neurons_=131, n_rats_=6). The vertical blue shaded bar represents the drug injection window. The slope graph (right) shows per-unit paired firing-rate changes; units with decreasing firing rate were highlighted in red. (B) Preferred theta phase (left) and phase locking values (right) before vs. after DA. DA 500 nmol induces spike-LFP theta synchrony and phase shift. The slope graph (right) shows per-unit paired phase locking values changes; units with increasing phase locking values were highlighted in red. (C-D) Saline condition shows no significant change (n_neurons_=70, n_rats_=6). # p<0.05 and ## p<0.01 for paired two-tailed t-tests.

### DA agonists SKF (D_1_) and QUIN (D_2_) do not reproduce DA-induced effects on HPC-PFC oscillatory dynamics

We next tested whether selective activation of D1- or D2-like receptors could reproduce the DA effects on HPC-PFC oscillatory activity at multiple doses. At theta frequencies, the selective D1 agonist SKF-38393 (1 or 10 µg) was able to modulate coherence (SKF vs. Sal - main effect: F_(2, 20)_=4.64, p=0.02, n_rats_=7; Fig. 5A, B, right), but not phase synchrony. However, this effect did not remain significant after adjustment (Holm-adjusted: p>0.05), suggesting that any differences were inconsistent across doses and not reliably different from saline. Overall, SKF was associated with non-significant modulation of theta relative power and HPC-PFC theta synchrony.

**Figure 5.**
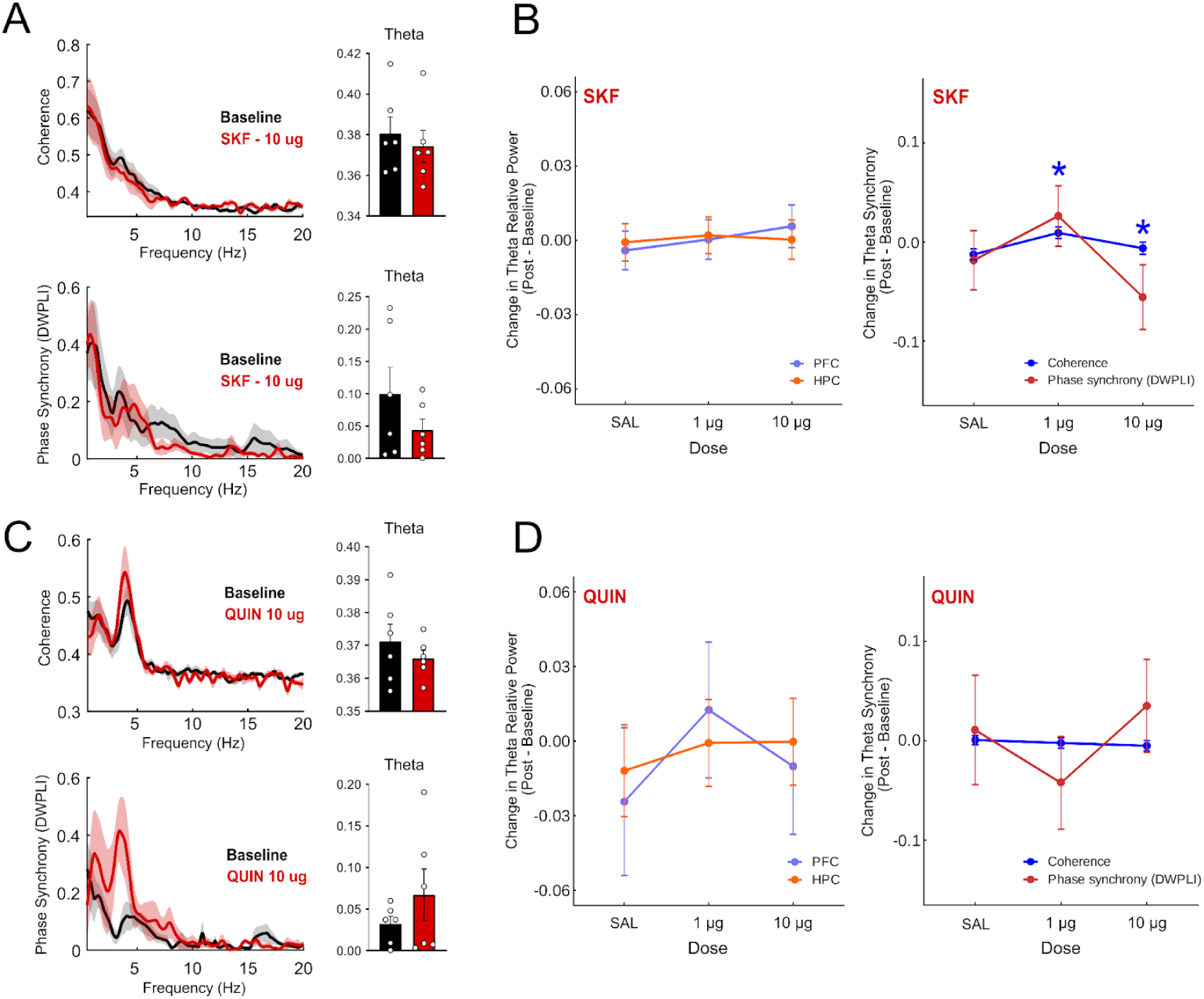
Selective D1 (SKF-38393) or D2 (quinpirole) receptor agonism does not reproduce DA-induced HPC-PFC theta modulation. (A) HPC-PFC coherence (top) and phase synchrony (dwPLI; bottom) averaged over 10-min post-injection vs. baseline, with theta bar plots (right; error bars represent SEM), for SKF 10 μg (n_rats_=7). (B) Dose-response of changes in theta relative power (left) and synchrony (right; coherence, dwPLI) across saline and both SKF doses (1 μg and 10 μg). (C) Coherence and phase synchrony for QUIN 10 μg with corresponding bar plots (n_rats_=6). Neither agonist produced consistent changes in theta power or synchrony; paired two-tailed t-tests. (D) Dose-response of power and synchrony across saline, and both QUIN doses (1 μg and 10 μg). For dose-response panels, points and error bars are linear mixed models (LMM) estimated marginal means ± SEM. * p<0.05 for LMM main effect.

The selective D2 agonist quinpirole (QUIN, 1 or 10 µg) similarly failed to induce reliable changes in HPC-PFC oscillatory activity. No consistent impact on theta relative power and on interareal synchrony measures was found, indicating no reliable effects produced by QUIN (Fig. 5C, D).

In contrast to DA, which increased theta power and HPC-PFC synchrony, isolated stimulation of D_1_ or D_2_ did not amplify theta relative power or strengthen interareal synchrony. This may indicate that concurrent D_1_/D_2_ activation is required to reproduce the DA-induced effects.

### Apomorphine exhibits a linear modulation of HPC-PFC oscillatory dynamics

To further investigate whether DA-induced effects can be reproduced by concurrent D1/D2 activation, we administered three different doses of the non-selective DA receptor agonist apomorphine (APO; 0.75, 1.5, and 3 mg/kg; i.p.). Due to APO systemic administration, analyses were conducted by contrasting the 30-40 min post-injection window with the injection baseline.

We found that APO affects HPC-PFC oscillatory activity in a dose-dependent manner. In fact, there was a main effect for theta power reduction on PFC (APO vs. Sal - main effect: F_(3, 22)_=3.76, p=0.03, n_rats_=7), but not HPC (Fig. 6B). However, this effect was not significant after post hoc adjustment (Holm-adjusted: p>0.05). At the lowest dose (0.75 mg/kg), the theta/delta relative power ratio indicated a theta-suppressing activity over time (Fig. 6A). At the intermediate dose (1.5 mg/kg), the most notable change was a predominance of theta activity over delta frequencies (Fig. 6A), which contrasted with the APO lowest dose. At the highest dose (3 mg/kg), there was a prominent inversion in the theta/delta relative power ratio compared to the lowest dose, with theta activity becoming predominant over time (Fig. 6A). Measures of inter-regional synchrony reflect the modulatory capacity of the APO, although the effects are not evident by comparing post-drug states directly with baseline levels. (Fig. 6C).

**Figure 6.**
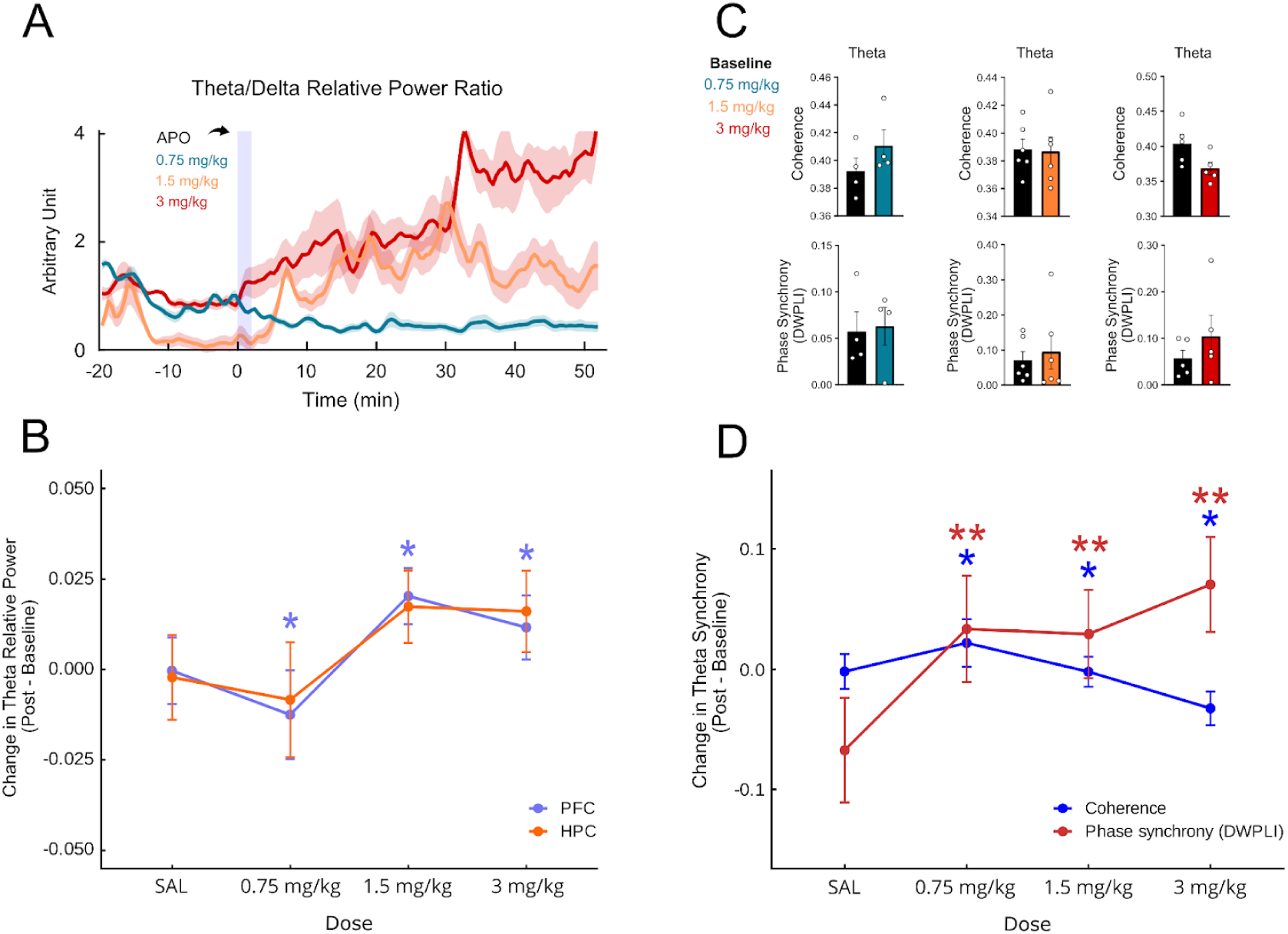
Apomorphine (i.p.) induces a dose-dependent modulation of HPC-PFC theta activity. (A) PFC theta/delta relative power ratio over time for 0.75, 1.5, and 3 mg/kg (normalized to baseline). Increasing the dose leads to an increase in theta relative power predominance over delta. The analysis window is 30-40 min post-dose to account for delays in the onset of effect following i.p. administration. Shaded areas around the lines represent SEM. (B) Dose-response of theta relative power change (post-injection - baseline). The magnitude of the change in relative power resembles the pattern shown by the theta/delta ratio. However, LMM analysis shows that the effect is more pronounced in the PFC. (C) Theta coherence (top row) and phase synchrony (bottom row) at each dose for post-injection vs. baseline, illustrating nonsignificant dose-linked shifts in synchrony measures when compared to baseline. Paired two-tailed t-tests. Error bars represent SEM. (D) Dose-response of theta synchrony (coherence, dwPLI). Relative to the saline injection, APO induces a strong positive modulation of theta phase synchrony while reducing theta coherence as the dose increases. For dose-response panels, points and error bars are linear mixed model (LMM) estimated marginal means ± SEM. For all plots, n_rats_=4 (0.75 mg/kg); n_rats_=6 (1.5 mg/kg); n_rats_=5 (3.0 mg/kg). * p<0.05 for LMM main effect; ** p<0.05 for LMM Holm-adjusted post hoc pairwise comparison.

APO produced a significant main effect on both coherence (APO vs. Sal: F(_3, ≈16_) = 4.06, p=0.03) and phase synchrony in the theta band (APO vs. Sal: F(_3, ≈14_) = 5.07, p=0.01). However, post hoc analyses revealed that only phase synchrony showed significant dose-dependent differences relative to saline, surviving Holm correction (0.75 mg/kg: p=0.04; 1.5 mg/kg: p=0.04; 3 mg/kg: p=0.02; Fig. 6D), whereas no individual dose reached statistical significance for coherence after correction. Together, these findings point to a predominant influence of APO on phase synchrony over amplitude-related coupling. Dose-response curves illustrate the dose-dependent action of APO, showing a modest modulation of relative power and coherence but a strong positive modulation of phase synchrony across theta frequencies as the dose increases (Fig. 6B, D).

## Discussion

In this study, we demonstrate that DA activation modulates HPC-PFC oscillatory dynamics under urethane anesthesia in a dose-dependent manner. Specifically, DA increases functional theta connectivity, enhances theta phase synchrony, and shifts PFC spike timing. Similarly, APO modulates the oscillatory activity. However, like the selective D_1_ and D_2_ receptor agonists it fails to reproduce the full spectrum of DA-induced effects.

Urethane anesthesia produces alternating activated and deactivated oscillatory states, whose activity patterns are similar to those observed in natural sleep states (Clement et al., 2008). During activated states, rapid low-amplitude theta oscillations and HPC-to-PFC 4 Hz synchronization predominate. In deactivated states, there are high-amplitude slow delta oscillations and a PFC to HPC 1 Hz synchronization predominance (Ruggiero et al., 2018; Lopes-Aguiar et al., 2020). Experimental studies have shown that sleep-wake cycles (Oishi & Lazarus, 2017) and urethane anesthesia are both regulated by D_1_- and D_2_-like dopaminergic receptors (Monti & Jantos, 2008). Our results demonstrate that DA and the non-specific agonist apomorphine, but not the selective D_1_ or D_2_ agonists, elicit dose-dependent alterations in urethane-induced oscillatory activity.

Both DA 100 nmol and 500 nmol doses induced theta oscillations predominance for at least 10 continuous minutes, but DA 500 nmol effects were more pronounced, showing higher and longer theta predominance. These results confirm previous studies, indicating active DA participation in HPC-PFC theta modulation (Benchenane et al., 2010; Ruggiero et al., 2021) and in sleep-wake-like systems (Oishi & Lazarus, 2017; Monti & Jantos, 2008). Interestingly, it was shown that acute DA depletion abolishes REM sleep, and REM can be recovered by D_2_ receptor stimulation (Dzirasa et al., 2006). However, D_2_-pathway activation is necessary but not sufficient by itself to generate REM sleep, and hyperdopaminergia is necessary but not sufficient to produce REM-like oscillations during wake in behaving DAT-KO mice (Dzirasa et al., 2006). This result indicates that the regulation of oscillatory states may also depend on the DA interaction with other neurotransmitter systems, such as the cholinergic system (Lester et al., 2010; Monti & Jantos, 2008). Indeed, cholinergic agonists can produce activated states, while cholinergic antagonists produce prolonged deactivated states (Clement et al., 2008; Ruggiero et al., 2018; Schall & Dickson, 2010). These results support the notion that dopaminergic and cholinergic systems may interact in regulating urethane-induced oscillatory states and the sleep-wake-like system (Dzirasa et al., 2006; Monti & Jantos, 2008).

We show that non-specific D_1_/D_2_ APO agonism has a dose-dependent effect on HPC-PFC oscillatory dynamics. In low doses, APO is known to produce sedative effects and increase the slow oscillations (Kropf & Kuschinsky, 1991; Monti & Jantos, 2008; Xu et al., 2011); high doses induce the opposite effect (Monti & Jantos, 2008). Our data confirm that low dose favored slower frequencies in the delta band and reduced theta predominance, while high dose induced theta-dominant activity. This effect is likely mediated by differential receptor mechanisms, with low doses preferentially activating dopaminergic autoreceptors, inhibiting neuronal firing, and decreasing DA release, whereas high doses stimulate postsynaptic receptors and enhance dopaminergic signaling (Monti & Jantos, 2008; Skirboll et al., 1979). Additionally, low APO doses reduce dopaminergic firing rate (Assié et al., 2009; Bunney et al., 1973a; 1973b; Mallet et al., 2008; Xu et al., 2011), whereas higher doses can promote neuronal excitability (Yanagihashi et al., 1991). At the highest doses, APO likely produced a hyperdopaminergic state. Still, APO is reported to be D_2_-biased (Dépatie & Lal, 2001), which may explain its partial divergence from DA in shaping HPC-PFC oscillations in our study. However, APO’s behavioral impact depends on intact D1 receptors (Ralph-Williams et al., 2002), highlighting that concurrent D1/D2 agonism can have effects that depend on which receptors are available.

We found that DA induced HPC-PFC theta synchrony and a phase shift in PFC neuronal activity. Consistent with this result, Benchenane et al. (2010) showed that DA infused into the PFC of anesthetized rats could induce HPC-PFC theta coherence and reorganize the preferred firing phase of PFC neurons relative to local theta cycles. However, to investigate HPC-PFC synchrony, Benchenane et al. (2010) relied solely on spectral coherence, which is biased by spectral power. The authors note that despite the increase in coherence, there is no alteration in spectral power in both HPC and PFC, suggesting that DA may induce phase synchrony independently of spectral power. By using dwPLI, a measure that quantifies consistent delays between signals and reduces zero-lag contamination due to volume conduction (Vinck et al., 2011), we showed that DA induced genuine HPC-PFC theta synchrony rather than just amplifying shared sources, and this effect was consistent across many animals following DA administration.

In our study, single-activation of D_1_ or D_2_ receptors by respective selective agonists was unable to induce significant effects. One possibility is that full dynamic dopaminergic modulation of HPC-PFC oscillatory synchrony requires concurrent activation of both D₁ and D₂ receptors to ensure a synergistic effect that no single-receptor agonist could achieve. For instance, this activation is required by cortical D₁-D₂ receptor heteromers to drive specific cellular and synaptic responses (Xu & Yao, 2010; Thompson et al., 2016; Perreault et al., 2014). However, the literature shows the involvement of specific dopaminergic receptors in modulating HPC-PFC activity. Xu et al. (2016) demonstrated in anesthetized rats that the administration of a D_1_ receptor antagonist could induce a reduction in HPC-PFC theta phase synchrony and theta power. Additionally, systemic administration of a D_1_ receptor agonist induced HPC-PFC theta coherence in pharmacological animal models of schizophrenia, but not in control animals (Perreault et al., 2017). In another study, a reduction in HPC-PFC theta phase synchrony was observed in freely-moving animals after systemic D_2_ receptor agonist or antagonist infusion (Gener et al., 2019). A likely explanation for why our results diverge from the literature regarding D_1_ and D_2_ effects is that the doses we used for selective agonists were suboptimal, and could mean insufficient receptor engagement at the low end, or activation of counter-regulatory mechanisms at the high end. Despite Perreault et al. (2017) and Gener et al. (2019) administering comparable agonist doses, it was delivered systemically, whereas we used the i.c.v. route, which may be related to the divergent findings.

More broadly, it is known that prefrontal DA mostly exerts non-linear, inverted U-shaped effects on cortical function, such that both hypo- and hyperdopaminergic states are associated with impaired cognition and only an intermediate range of receptor stimulation is optimal (Weber et al., 2022), highlighting the complex and dose-sensitive nature of dopaminergic modulation. The complex DA receptor pharmacodynamics, including differential activation of D_1_ and D_2_ receptors, autoreceptors, and D_1_-D_2_ heteromers across dose ranges, further complicate the interpretation of our negative findings with selective agonists. Also, a reduced sample size in the agonist treatment groups may explain the lack of statistical significance, an issue partially mitigated by our linear mixed-effects models, which account for repeated measures and between-animal variability. Future work using broader dose-response designs, receptor-specific antagonism, and cell-type or projection-specific manipulations will be essential to learn how these receptor mechanisms shape HPC-PFC synchrony. Importantly, our results were obtained under anesthesia, and it is known that urethane anesthesia neuromodulatory tone and state transitions differ from wakefulness. Thus, follow-up experiments in behaving animals are needed to generalize the results found in this study, and to better understand DA’s role in interareal synchronization across wakefulness states.

Our findings link DA to long-range circuit-level communication, showing that DA can rapidly increase low-frequency synchrony in crucial cognitive networks (HPC-PFC), beyond simply changing local activity. In Parkinson’s disease, cognitive impairment has been associated with reduced midfrontal low-frequency rhythms, motivating the investigation of biomarkers and stimulation-based approaches to restore coordination (Singh et al., 2021; Narayanan et al., 2024). Our results suggest that DA loss may weaken not only low-frequency frontal oscillations but also long-range HPC-PFC coupling needed for cognitive performance. Clinically, this circuit-level interpretation aligns with the use of apomorphine in patient cohorts reporting improvements in aspects of non-motor symptom burden, including in patients with cognitive and neuropsychiatric dysfunctions (Carbone et al., 2019). In schizophrenia, DA dysregulation and network dysconnectivity are accompanied by medial frontal low-frequency deficits during cognitive timing (Parker et al., 2017). Also, in an amphetamine hyperdopaminergic model, an increase in timing variability and disruption of prefrontal temporal coding has been shown while also reducing low-frequency oscillations (Weber et al., 2025). Together, these observations support the translational idea that DA may help to regulate cognition by tuning the timing of interactions across circuits, making low-frequency coupling and spike timing potential targets for pharmacology and neuromodulation in DA-related brain disorders.

## Conclusion

We demonstrate that DA dose-dependently reconfigures HPC-PFC network dynamics under urethane anesthesia, increasing theta power, enhancing theta phase synchrony, and strengthening prefrontal spike-theta locking. In contrast, selective D_1_ or D_2_ agonism failed to reproduce these effects, and apomorphine produced a dose-dependent modulation of oscillatory dynamics. Importantly, phase-coupling analyses that minimize zero-lag effects, thereby reducing volume-conduction and power-related confounds, support the interpretation that the observed synchrony reflects genuine interareal coupling, suggesting that DA rapidly regulates hippocampal-prefrontal communication.

## Acknowledgments

We thank Antonio Renato Meirelles e Silva, Renata Caldo Scandiuzzi, and Daniela Ribeiro for their technical support, and Tamiris Prizon, José Luiz Liberato, and Leonardo Rakauskas Zacharias for their experimental support. The research was conceptualized by RNR and BAOJ. BAOJ conducted the experiments. BAOJ and RNR analyzed data. BAO-J, RNR, FEP-N, and NSN wrote the manuscript. RNR and JPL supervised the research.

